# Presaccadic attention enhances and reshapes the Contrast Sensitivity Function differentially around the visual field

**DOI:** 10.1101/2023.11.16.567379

**Authors:** Y. Kwak, Y. Zhao, Z-L. Lu, N.M. Hanning, M. Carrasco

## Abstract

Contrast sensitivity, which constrains our vision, decreases from fovea to periphery, from the horizontal to the vertical meridian, and from the lower vertical to the upper vertical meridian. The Contrast Sensitivity Function (CSF) depicts how contrast sensitivity varies with spatial frequency (SF). To overcome these visual constraints, we constantly make saccadic eye movements to foveate on relevant objects in the scene. Already before saccade onset, *presaccadic attention* shifts to the saccade target and enhances perception. However, it is unknown whether and how it modulates the interplay between contrast sensitivity and SF, and if this effect varies around polar angle locations. Contrast sensitivity enhancement may result from a horizontal or vertical shift of the CSF, increase in bandwidth, or any combination. Here, we investigated these possibilities by extracting key attributes of the CSF using Hierarchical Bayesian Modeling, which enables precise estimation of the CSF parameters by decomposing the variability of the dataset into multiple levels. The results reveal that presaccadic attention (1) enhances contrast sensitivity across SF, (2) increases the most preferred and the highest discernable SF, and (3) narrows the bandwidth. Therefore, presaccadic attention bridges the gap between presaccadic and post-saccadic input by increasing visibility at the saccade target. Counterintuitively, the presaccadic enhancement in contrast sensitivity was more pronounced where perception is better –along the horizontal than the vertical meridian– exacerbating polar angle asymmetries. Our results call for an investigation of the differential neural modulations underlying presaccadic perceptual changes for different saccade directions.

**Significance statement:** The contrast sensitivity function (CSF) describes how our ability to perceive contrast depends on spatial frequency. Contrast sensitivity is highest at the fovea and decreases in the periphery, especially at vertical locations. We thus make saccadic eye movements to view objects in detail. Already before moving our eyes, *presaccadic attention* enhances perception at the target location. But how does it influence the interplay between spatial frequency and contrast sensitivity, and does its effect vary around the visual field? Using Hierarchical Bayesian Modeling, we show that presaccadic attention enhances and reshapes the CSF to prepare the periphery for upcoming fixation. Counterintuitively, it does so more at horizontal locations where vision is stronger, suggesting smoother perception across horizontal than vertical eye movements.

## 1. Introduction

Visual processing varies across eccentricity and around polar angle locations. Our ability to process fine details at the center of gaze declines with eccentricity (Frisén & Glansholm, 1975). Moreover, at iso-eccentric locations, visual performance changes as a function of polar angle (**Figure 1a**, review: Himmelberg, Winawer, & Carrasco, 2023): it is better along the horizontal than vertical meridian (HVA: Horizontal-Vertical Anisotropy) and along the lower than the upper vertical meridian (VMA: Vertical Meridian Asymmetry).

**Figure 1.**
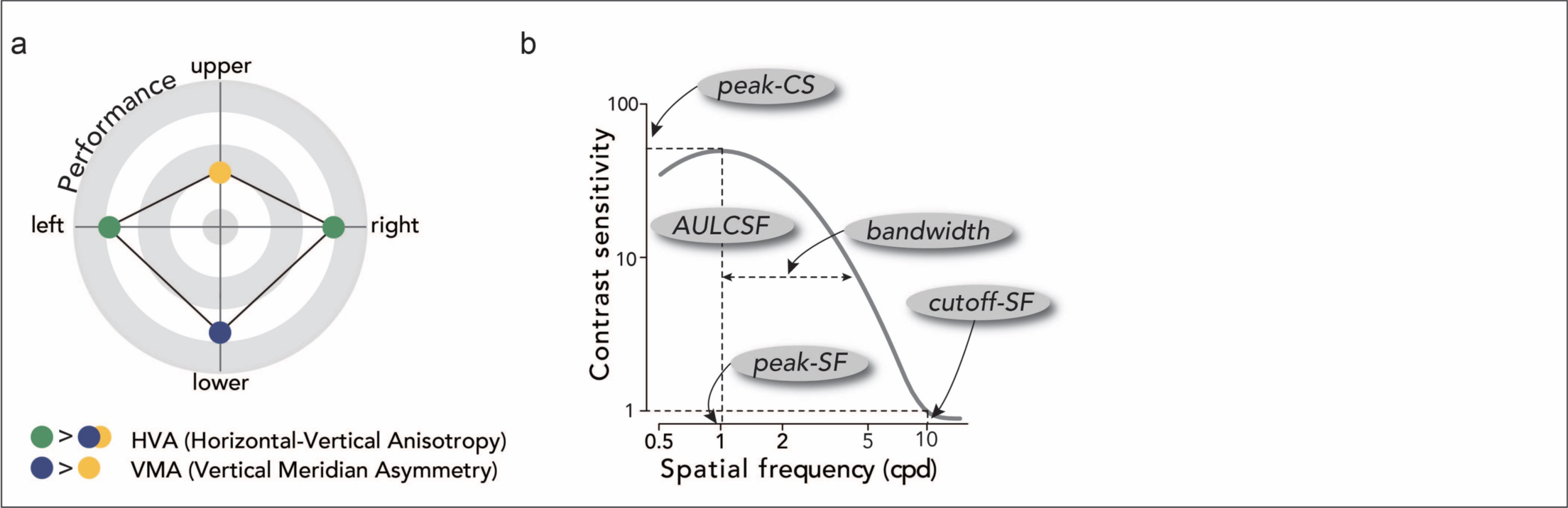
Background. **(a)** Polar angle asymmetries around the visual field. Visual performance is better along the horizontal than the vertical meridian, and along the lower than the upper vertical meridian. **(b)** Key attributes of the CSF. We derived peak-CS (peak contrast sensitivity), peak-SF (peak spatial frequency), bandwidth (full width at half maximum; half width shown for illustration purposes), cutoff-SF (cutoff spatial frequency), and AULCSF (area under the log CSF). Key CSF attributes were measured during fixation and saccade preparation to quantify presaccadic attention modulation on the CSFs.

We often compensate for these visual constraints by making rapid saccadic eye movements to actively explore the world and place relevant objects in the fovea where visual perception is most sensitive (Eckstein, 2017). Already before our eyes move (during saccade preparation), *presaccadic attention* shifts to the saccade target location and prioritizes visual processing (e.g., Deubel & Schneider, 1996; Kowler, Anderson, Dosher, & Blaser, 1995; Montagnini & Castet, 2007), Presaccadic attention boosts accuracy (e.g., Hanning, Deubel, & Szinte, 2019; Hoffman & Subramaniam, 1995) and reduces reaction times (e.g., Shepherd, Findlay, & Hockey, 1986). Some studies investigating how presaccadic attention modulates perception of basic visual dimensions have revealed that saccade preparation enhances contrast sensitivity (Hanning, Fernández, & Carrasco, 2023; Hanning, Himmelberg, & Carrasco, 2022, 2023; Li, Pan, & Carrasco, 2021) and spatial acuity (Kwak, Hanning, & Carrasco, 2023); sharpens orientation tuning and orientation acuity (Castet, Jeanjean, Montagnini, Laugier, & Masson, 2006; Li, Barbot, & Carrasco, 2016; Ohl, Kuper, & Rolfs, 2017); and shifts spatial frequency (SF) tuning by enhancing sensitivity to higher SFs (Li et al., 2016, 2019). The presaccadic attention effect increases as saccade onset approaches (Hanning et al., 2019; Kroell & Rolfs, 2021; Li et al., 2016; Ohl et al., 2017; Rolfs & Carrasco, 2012). These studies suggest a functional role of presaccadic attention in rendering visual information at the peripheral saccade target more “fovea-like”. Across every saccade, two distinct percepts need to be seamlessly integrated: a blurry presaccadic input from the peripheral saccade target and a high-resolution post-saccadic input from the same location once the saccade target has been foveated. Presaccadic attention eases the integration of pre- and post-saccadic visual information, facilitating perceptual continuity (Herwig, 2015; Herwig & Schneider, 2014; Kroell & Rolfs, 2022; Li, Hanning, & Carrasco, 2021; Stewart, Valsecchi, & Schütz, 2020).

Contrast sensitivity constrains our vision. It drops markedly with eccentricity (Rovamo, Virsu, & Näsänen, 1978; Virsu & Rovamo, 1979) and exhibits pronounced polar angle asymmetries (Abrams, Nizam, & Carrasco, 2012; Baldwin, Meese, & Baker, 2012; Jigo, Tavdy, Himmelberg, & Carrasco, 2023; Rijsdijk, Kroon, & van der Wildt, 1980). Moreover, contrast sensitivity depends heavily on SF and is bandpass (**Figure 1b**). At the fovea, for example, it peaks at a SF range of 2 to 6cpd and declines rapidly for lower and higher SFs (Campbell & Robson, 1968; Kelly, 1977). This general relation between contrast sensitivity and SF, referred to as the Contrast Sensitivity Function (CSF), defines our “window of visibility” (Ginsburg, 1978; Watson, Ahumada, & Farrell, 1986).

Presaccadic attention has been shown to increase contrast sensitivity across the contrast response function *at a constant SF* (Hanning et al., 2022; Li, Pan, et al., 2021; Zhao, Gersch, Schnitzer, Dosher, & Kowler, 2012), and performance at a range of SFs at a *constant contrast* (Kroell & Rolfs, 2021). But how It modulates the interplay between contrast and SF sensitivity is unknown: An increase in contrast sensitivity at a given SF may result from an upward shift of the CSF, a horizontal shift, increased bandwidth, or any of their combinations (**Figure 1b**).

Here, we investigated presaccadic modulations jointly by measuring the CSF around the visual field. To investigate whether and how presaccadic attention shapes our window of visibility, we characterized the CSFs and compared the key CSF attributes (**Figure 1b**) during fixation and saccade preparation. Furthermore, as polar angle asymmetries established during fixation (**Figure 1a**) are exacerbated during saccade preparation for contrast sensitivity (Hanning et al., 2022, 2023) but are preserved for acuity (Kwak et al., 2023), here we evaluated presaccadic modulations of the CSF as a function of saccade direction. We fit a Hierarchical Bayesian Model (HBM) that has been applied to analyze CSFs (Zhao, Lesmes, Dorr, & Lu, 2021; Zhao, Lesmes, Hou, & Lu, 2021). The HBM decomposes the variability of the dataset into multiple hierarchies and takes into consideration potential relations between CSF parameters. Thus, the model enables precise estimates of the peak contrast sensitivity (peak-CS), the preferred SF at the peak-CS (peak-SF), bandwidth (full width at half maximum), the highest discernable SF (cutoff-SF), and area under the log CSF (AULCSF).

We found polar angle asymmetries in peak-CS, cutoff-SF, and AULCSF during fixation (Jigo et al., 2023). Importantly, presaccadic attention 1) shifted the CSF upward, increasing the peak-CS; 2) shifted the CSF rightward to higher SFs, increasing the peak-SF and cutoff-SF; 3) narrowed the CSF, decreasing the bandwidth; and 4) increased the AULCSF. These presaccadic modulations were present at all polar angle locations, but there was a larger presaccadic enhancement along the horizontal than the vertical meridian, specifically for the peak-CS as well as for contrast sensitivity across a wide range of SFs. These findings reveal that presaccadic attention increases our window of visibility, reducing the gap between visual representations at the peripheral saccade target and the future fovea, and more so before horizontal than vertical saccades.

## 2. Materials and Methods

### 2.1. Participants

Twelve observers (4 males and 8 females, including authors YK and NH; ages 20–34) with normal or corrected-to-normal vision participated in the experiment. All participants (except for the two authors) were naïve to the experimental hypothesis and were paid $12 per hour. The experimental procedures were approved by the Institutional Review Board at New York University, and all participants (except for the two authors) provided informed consent. All procedures were in agreement with the Declaration of Helsinki.

### 2.2. Setup

Participants sat in a dark room with their head stabilized by a chin and forehead rest. All stimuli were generated and presented using MATLAB (MathWorks, Natick, MA, USA) and the Psychophysics Toolbox (Brainard, 1997; Pelli, 1997) on a gamma-linearized 20-inch ViewSonic G220fb CRT screen (Brea, CA, USA) at a viewing distance of 79 cm. The CRT screen had a resolution of 1,280 by 960 pixels and a refresh rate of 100 Hz. Gaze position was recorded using an EyeLink 1000 Desktop Mount eye tracker (SR Research, Osgoode, Ontario, Canada) at a sampling rate of 1 kHz. The Eyelink Toolbox was used for eyetracking with MATLAB and Psychophysics Toolbox (Cornelissen, Peters, & Palmer, 2002).

### 2.3. Experimental Procedure

Participants performed an orientation discrimination task for stimuli varying in contrast and spatial frequency, during *fixation* and *saccade preparation (presaccadic attention)* (**Figure 2**). To place trials efficiently in the dynamic range of the CSF, we leveraged the qCSF (quick Contrast Sensitivity Function) method for data collection (Lesmes, Lu, Baek, & Albright, 2010). Given participants’ performance in previous trials, this procedure selects stimulus contrast and spatial frequency to be tested on each trial, based on the maximum expected information gain, to further refine the CSF parameter estimates. The possible stimulus space was composed of 60 contrast levels from 0.001 to 1, and 12 spatial frequency levels from 0.5 to 16cpd, evenly spaced in log units. We manipulated eye movement instruction (*fixation* and *saccade preparation,* from now on “instruction”) between experimental blocks. **Figure 2a** shows the trial sequence under the saccade instruction. Each trial started with a fixation circle (0.175° radius) on a gray background (∼26 cd/m^2^) with a duration randomly jittered between 400ms and 600ms. Four placeholders indicated the locations of the upcoming stimuli, 6° left, right, above, and below fixation. Measurements at the left and right locations were combined for analysis, as it is well established that contrast sensitivity does not differ between these locations (e.g., Cameron, Tai, & Carrasco, 2002; Himmelberg, Winawer, & Carrasco, 2020). Each placeholder was composed of four corners (black lines, 0.2° length). The trial began once a 300-ms stable fixation (eye coordinates within a 1.75° radius virtual circle centered on fixation) was detected.

**Figure 2.**
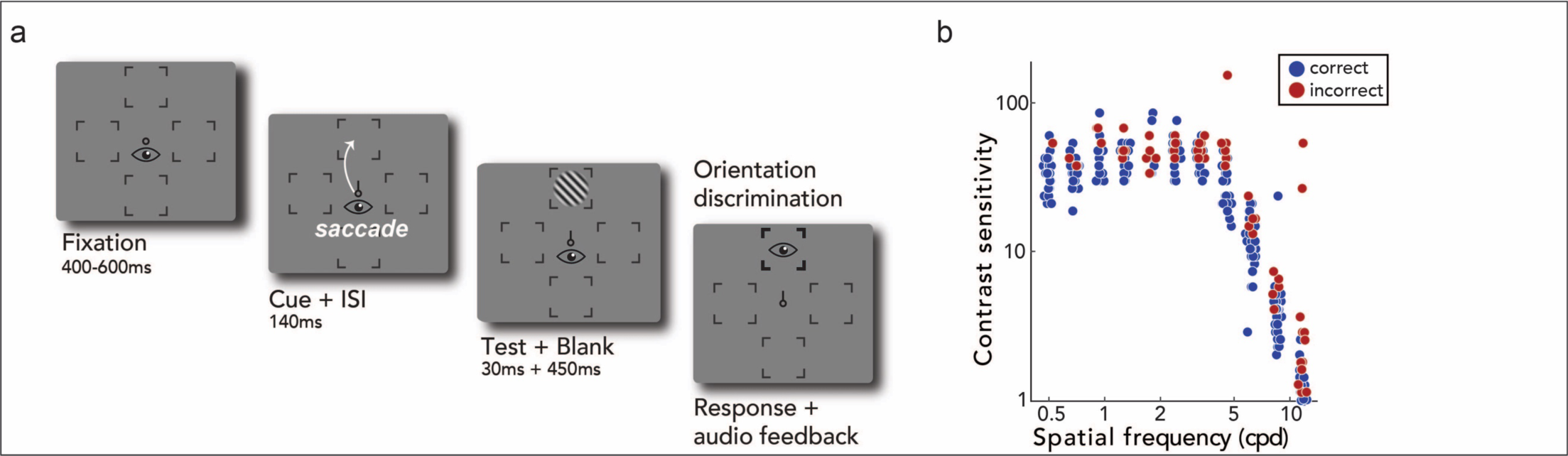
Experimental procedure. (a) Trial sequence. In saccade blocks (*presaccadic attention*), a central saccade cue indicated the saccade target after a fixation period. Participants were instructed to make a saccade precisely to the center of the corresponding placeholder as soon as the cue appeared. Just before saccade onset, a test Gabor stimulus was presented briefly at the cued location, and participants judged its orientation (CW or CCW) after response cue onset (i.e., after the saccade). The spatial frequency and the contrast of the Gabor on each trial was varied based on the qCSF (quick Contrast Sensitivity Function) procedure. The eye icons are for illustration only and were not shown during the experiment. In the fixation blocks (not illustrated here), the trial sequence was identical except that the cue pointed to all four locations, and participants were instructed to fixate at the center throughout the entire block. (b) Trial-by-trial data points for an example participant and location. Blue and red dots indicate trials with correct and incorrect responses respectively. The qCSF procedure efficiently samples stimuli in spatial frequency and contrast space based on responses on previous trials.

In blocks with the saccade instruction (*saccade preparation*), after the fixation period, a central cue (black line, 0.35° length) pointed to one of the four cardinal placeholders, indicating the saccade target. Participants were instructed to move their eyes to the center of the cued location as fast and precisely as possible. 140ms after cue onset (i.e., before saccade onset), a test Gabor grating (tilted ±45 relative to vertical, random phase, delimited by a raised cosine envelope with 2° radius) appeared for 30ms at the cued saccade target location. The spatial frequency and the contrast of the Gabor stimulus were determined by the qCSF procedure on each trial (Lesmes et al., 2010), based on the maximum expected information gain (reduction in entropy). 450ms after stimulus offset (and after saccade offset), the cued location was highlighted by increasing the width and length of the corresponding placeholders. Participants performed an un-speeded, orientation discrimination task for the Gabor grating at the cued location (by pressing left arrow key for -45°, right arrow key for +45°).

Stimulus parameters and timing for the blocks with the fixation instruction were identical to the saccade blocks with one exception: instead of one saccade target cue, four black direction cue lines pointed to all locations. Participants maintained fixation throughout the entire trial sequence.

Gaze position was monitored online to ensure fixation within a 1.75° radius virtual circle from the central fixation until the response phase (fixation blocks) or until cue onset (saccade blocks). Trials in which gaze deviated from fixation were aborted and repeated at the end of each block. In addition, in the saccade blocks, we repeated trials in which saccade latency was too short (< 150ms) or too long (> 350ms), or in which saccades landed outside of a 2.25° radius virtual circle around the saccade target center. Moreover, because we are interested in the presaccadic interval, we only included trials in which saccades were initiated after stimulus offset.

### 2.4. CSF analysis

#### 2.4.1. Bayesian Inference

To characterize the CSFs during *fixation* and *saccade preparation* for each of the four locations, we used the Bayesian Inference Procedure (BIP, **Figure 3a**) and the Hierarchical Bayesian Model (HBM, **Figure 3b**) to fit the three-parameter CSF model to trial-by-trial data points (**Figure 2b**) (Zhao, Lesmes, Dorr, et al., 2021; Zhao, Lesmes, Hou, et al., 2021). For both the BIP and the HBM, Bayesian inference was used to estimate the posterior distribution of the three CSF parameters Dpeak contrast sensitivity (peak-CS), peak spatial frequency (peak-SF), and bandwidth. Other key attributes, such as the cutoff spatial frequency (cutoff-SF) and area under the log CSF (AULCSF), were computed from the final estimate of the CSF.

**Figure 3.**
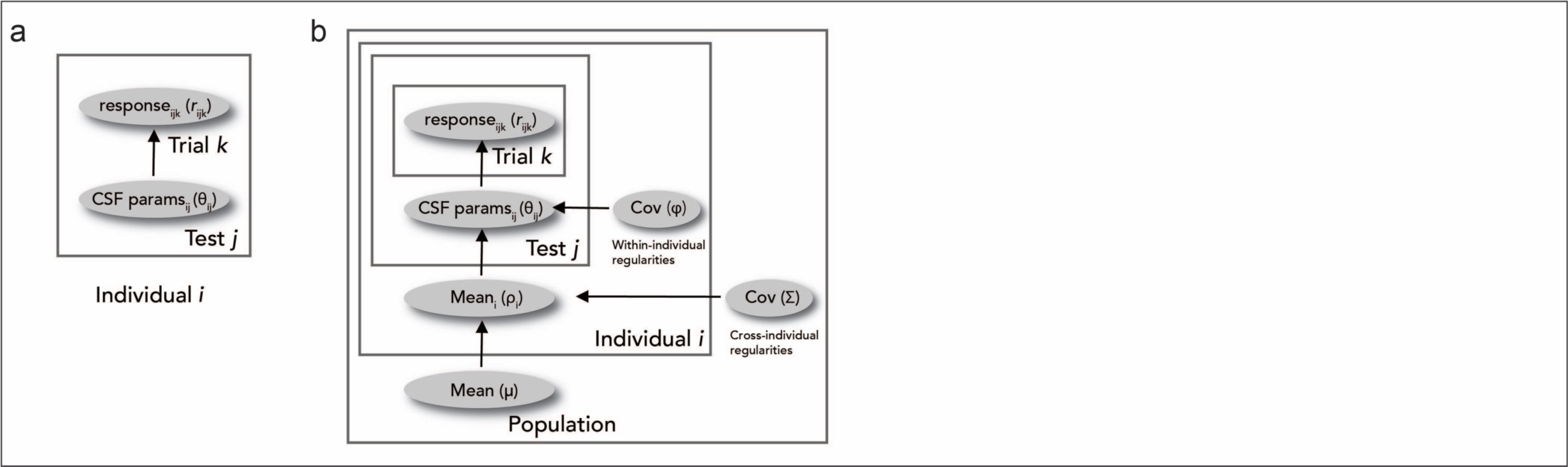
CSF model fitting, (a) Schematic representation of the BIP (Bayesan-terence Procedure). The fist soge of the data analysis consists of fitting trial-by-trial data ports uith the 81P to estm3te the CSF parameters —pe3k contrast sensitivity (peak-CS). peak spatial frequency (peak-SF). and bandwidth. The BIP computes the posterior distribution of the parameters for each test independently, (b) Schematic representation of the HBM (Hierarchical Bayesian Model). In the second stage of the data analysis, the BiP outputs are used as the mean and covananoe of the pnor dstnbutons of the CSF hyperparameters in the HBM. We used a three-level hierarchical model to incorporate potential relations in the CSF parameters across individuals and tests. For details on the BIP and the HBM. see 2.4. CSF analysis.

Contrast sensitivity *S* at spatial frequency *sf* is modeled as a log-parabola function with parameters θ = (*peakCS*, *peakSF*, *bandwidth*) (Lesmes et al., 2010; Watson & Ahumada, 2005):

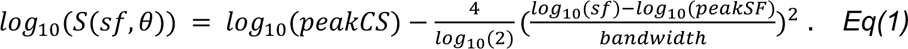

The probability of a correct response (*r* = 1) on a trial given the stimulus —spatial frequency *sf* and contrast *c* —is described as a psychometric function:

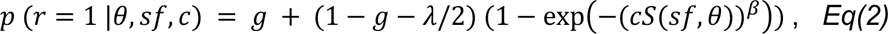

where *g* is the guessing rate (*g* = 0.5), λ is the lapse rate (λ = 0.04; Wichmann & Hill, 2001; Zhao, Lesmes, Hou, et al., 2021), and β determines the slope of the psychometric function (β = 2). The probability of making an incorrect response (*r* = 0) is:

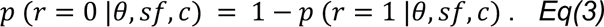

Equations 2 and 3 define the likelihood function, or the probability of a correct/incorrect response in a trial given the stimulus and the CSF parameters. To infer the CSF parameters given the experimental data, Bayes’ Rule is used to estimate the posterior distribution of the CSF parameters (θ) based on the likelihood and the prior (*Eq(7)*) in the BIP procedure. Details on modeling the prior distributions can be found in the **Supporting Information**.

#### 2.4.2. HBM: Three-level hierarchy

We used a simple three-level HBM without considering any structure related to the experimental conditions, as reported in Zhao, Lesmes, Dorr, et al. (2021) (**Figure 3b**). In the current study, an individual refers to a combination of a participant, location (upper, lower, horizontal), and instruction (*fixation*, *saccade preparation*) condition (*I*=72). In addition, all trials that each participant completed were combined into one test (*J*=1).

The BIP fits these parameters independently for each individual and test (**Figure 3a**). Although the BIP has been proven to be a good estimate of the CSF, it may have overestimated the variance of each test because it scores each test independently with a uniform prior without considering potential relations of the parameters (Zhao, Lesmes, Dorr, et al., 2021). Therefore, we used a hierarchical model which recently has been shown to reduce the uncertainties of the parameter estimates when fitting the CSF (Zhao, Lesmes, Dorr, et al., 2021; Zhao, Lesmes, Hou, et al., 2021). The HBM considers potential relations of the CSF parameters and hyperparameters within and across individuals (**Figure 3b**). More specifically, it quantifies the joint distribution of the CSF parameters and hyperparameters at three hierarchies in a single model: test-(*J*), individual-(*I*), and population-level. The within-individual and cross-individual regularities are modeled as the covariance of the CSF parameters at the individual level (φ) and covariance at the population level (Σ), respectively. Incorporating this knowledge into the model and decomposing the variability of the entire dataset into distributions at multiple hierarchies enabled us to reduce the variance of the test-level estimates and to obtain more precise estimates of the CSF parameters.

We fit the BIP and the HBM sequentially. As a first step, the BIP was fit to obtain the mean and standard deviation of each of the three CSF parameters across all participants and conditions, using a uniform prior distribution. Next, these values were fed into the HBM and set the mean and covariance of the prior distributions of the CSF hyperparameters (*Eq(S1)*).

At the population level of the HBM, the joint distribution of hyperparameter η was modeled as a mixture of three-dimensional Gaussian distributions *N* with mean Σ and covariance Σ, which have distributions *p*(μ) and *p*(Σ):

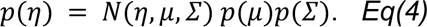

At the individual level (*I*), the joint distribution of hyperparameter τ_*i*_ of individual *i* was modeled as a mixture of three-dimensional Gaussian distributions *N* with mean ρ_*i*_ and covariance φ, which have distributions *p*(ρ_*i*_|η) and ρ(φ); where ρ(ρ_*i*_|η) denotes that the mean ρ_*i*_ was conditioned on the population-level hyperparameter η:

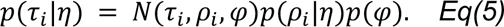

At the test level (*J*), *p*(θ_*i,j*_), the joint distribution of parameter θ_7;_ of individual *i* and test *j* was conditioned on hyperparameter τ_*i*_ at the individual level.

The probability of obtaining the entire dataset was computed by probability multiplication:

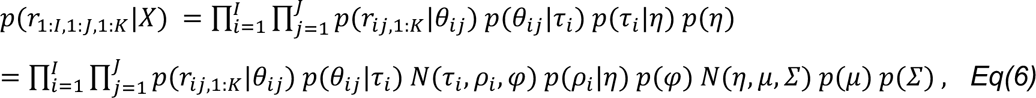

where *X* = (θ_1:*I*, 1:*J*_, ρ_1: *i*_ φ, μ, Σ) are all parameters and hyperparameters in the HBM, *I* is the total number of individuals, and *J* is the total number of tests on each individual.

#### 2.4.3. HBM: Computing the joint posterior distribution

Bayes’ rule 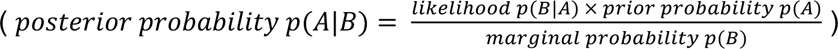 was used to compute the joint posterior distribution of all the parameters and hyperparameters in the HBM:

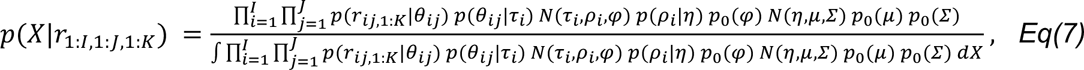

where the denominator is the probability of obtaining the entire dataset (*Eq(6)*).

We used the JAGS package (Plummer. 2003) in R (R Core Team. 2020, to evaluate the joint posterior distribution. JAGS generates representative samples of the joint posterior distribution of all the parameters and hyperparameters in the HBM via Markov Chain Monte Carlo (MCMC). We ran three parallel MCMC chains, each generating 2000 samples, resulting in a total of 6000 samples. Steps in the burn-in and adaptation phases — 20000 and 500000 steps respectively— were discarded 3nd excluded from the analysis because the initial part of the random walk process is largely determined by random starting values.

### 2.5. Eye movement analysis

In addition to the gaze monitoring and detection of saccades online (see **2.3. Experimental Procedure**), we also performed an offline eye movement analysis using an established algorithm for saccade detection (Engbert & Mergenthaler, 2006). Saccades were detected based on their velocity distribution using a moving average over 20 subsequent eye position samples. Saccade onsets and offsets were detected based on when the velocity exceeded or fell below the median of the moving average by 3 standard deviations for at least 20ms. This offline analysis takes into account the entire distribution of eye position samples and velocity and allows for a more detailed evaluation of both fixation and saccadic eye movements. In total, we included 22,527 trials in the analysis Don average 1877 ± 55 (mean ± 1 SEM) trials per observer.

### 2.6. Statistical analysis

Convergence of the HBM parameters was determined based on the Gelman and Rubin’s diagnostic rule (Gelman & Rubin, 1992). Each parameter and hyperparameter was considered to have “converged” when the variance of the samples across the MCMC chains divided by the variance of the samples within each chain was smaller than 1.05. In the current study, all parameters of the HBM converged.

To obtain final estimates of the CSF parameters in the HBM 〡peak-CS, peak-SF, and bandwidth〡, parameter estimates of the 6000 samples were averaged for each participant and location-instruction combination. The final CSFs for each participant and location-instruction combination were obtained by inputting the final parameter estimates to *Eq(1)*. From these final CSFs, cutoff-SF and AULCSF were computed. These key CSF attribute values were used for statistical testing.

All *p* values reported are based on permutation testing over 1000 iterations with shuffled data. *P* value is the proportion of a metric (*F* score or *t* score) in the permuted null distribution greater than or equal to the metric computed using intact data. Note that the minimum *p* value achievable with this procedure is 0.001. *P* values were FDR corrected for multiple comparisons, when applicable (Benjamini & Hochberg, 1995).

We also performed Bayesian statistics using the “BayesFactor” package in R (version 0.9.12–4.4). We report BF_10_ only for null results from permutation testing. The Bayes factors (BF_10_) for main effects and interaction effects in the repeated measures ANOVA design were computed by comparing the full model (H1) against the restricted model (H0) in which the effect of interest was excluded from the full model (Rouder, Morey, Verhagen, Swagman, & Wagenmakers, 2017). BF_10_ smaller than 1 is in support for the absence of an effect; values 1–3, 3–10, 10–30, 30–100, and > 100 indicate anecdotal, moderate, strong, very strong, and extreme evidence for the presence of an effect (Kass & Raftery, 1995; Lee & Wagenmakers, 2014).

## 3. Results

Figure 4 shows the CSFs averaged across participants, during *fixation* and *saccade preparation* (presaccadic attention) for each location. To preview the results, *saccade preparation* shifted and reshaped the CSFs during *fixation* for all locations, increasing overall contrast sensitivity (**Figure 4a**). Second, during *fixation* (**Figure 4b**), contrast sensitivity was better in the horizontal than the vertical meridian, and in the lower vertical than the upper vertical meridian, consistent with HVA and VMA effects. Third, this pattern was overall maintained during *saccade preparation* (**Figure 4c**), but the presaccadic benefit on contrast sensitivity was more pronounced at the horizontal than at the vertical meridian across a wide spatial frequency range.

**Figure 4.**
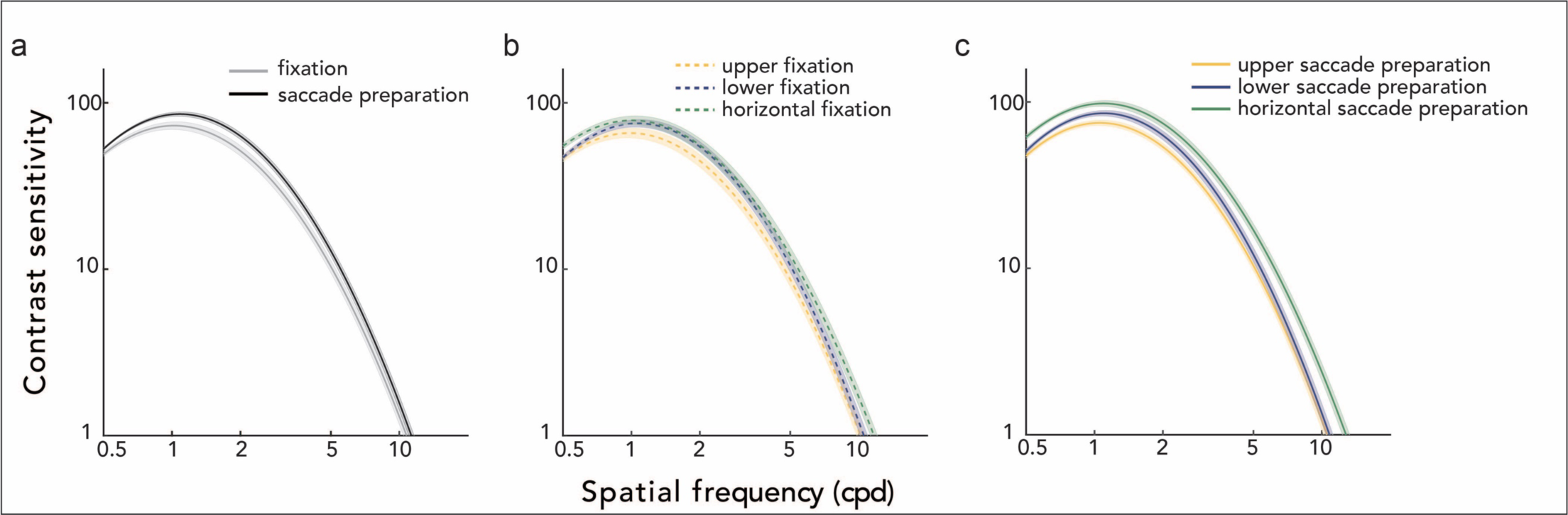
Group average CSFs. **(a)** CSFs for the *fixation* and *saccade preparation* instruction, collapsed across locations. *Saccade preparation* modulates contrast sensitivity across spatial frequencies. **(b)** CSFs during *fixation* for each location, exhibiting polar angle asymmetries. **(c)** CSFs during *saccade preparation*, separately for each location. Polar angle asymmetries in the CSFs during *fixation* were also present during *saccade preparation*. **(a-c)** The CSF parameters were fitted with the HBM (Hierarchical Bayesian Model). Shaded regions denote ±_1_ standard error of the mean (SEM).

We conducted a two-way repeated measures ANOVA for each of the key CSF attributes, with location (upper, lower, horizontal) and eye movement instruction (*fixation*, *saccade preparation*) as within-subject factors (**Figure 5**).

**Figure 5.**
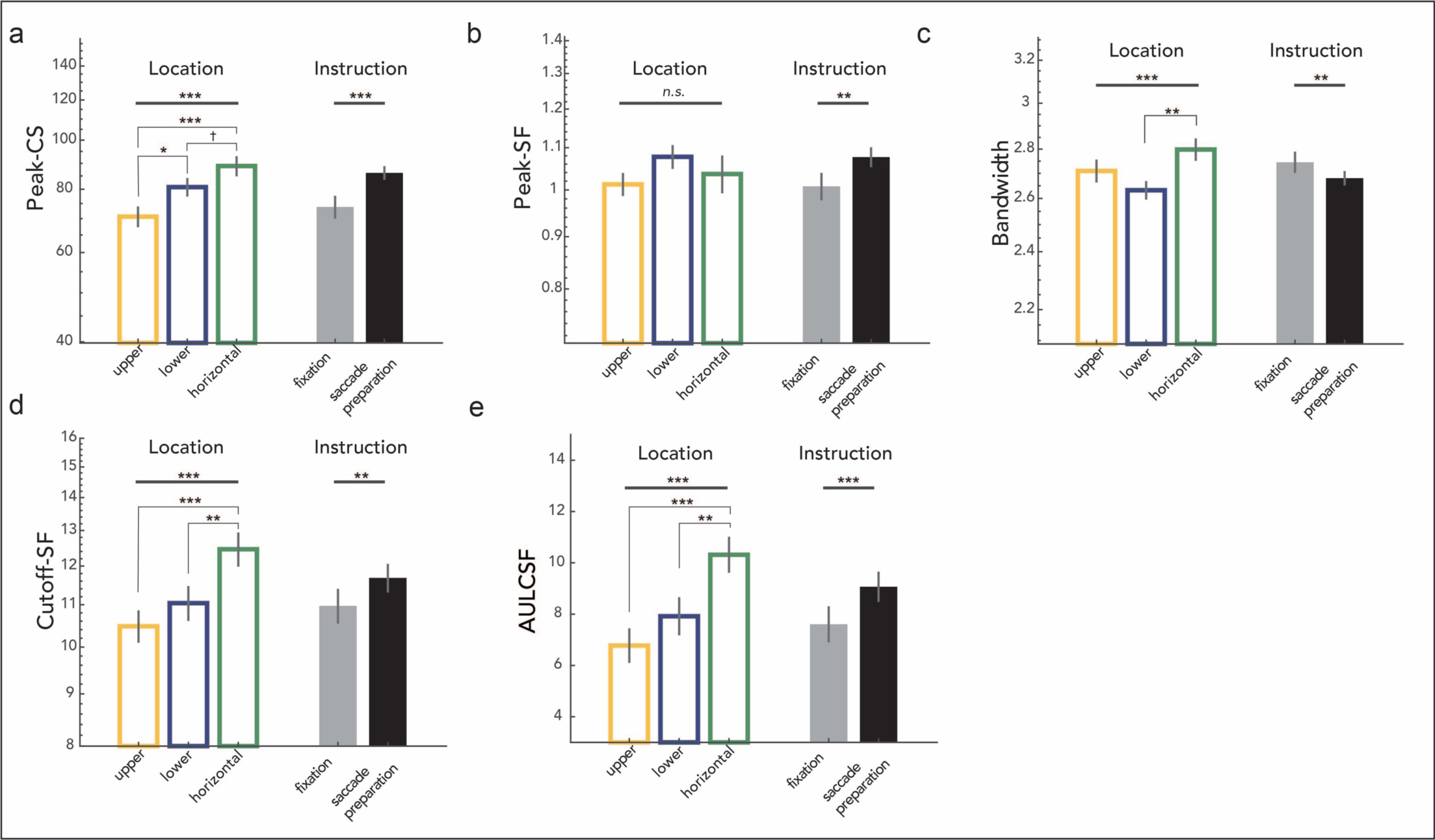
Main effects of location (averaged across *fixation* and *saccade preparation* instruction) and eye movement instruction (averaged across locations). **(a)** peak-CS; **(b)** peak-SF; **(c)** bandwidth; **(d)** cutoff-SF; **(e)** AULCSF. Errorbars indicate ±_1_ standard error of the mean (SEM), and vertical bars depict ±_1_ standard error of the difference (SED). Main effects of location and instruction are shown above the thick gray horizontal bars on the top. Significant post-hoc pairwise comparisons of location effects are denoted by asterisks above thin gray bars. \*\*\**p* < 0.001, \*\**p* < 0.01, \**p* < 0.05, †*p* < 0.1, *n.s.* not significant; FDR corrected.

### 3.1. Peak-CS: peak contrast sensitivity

We observed main effects of location (*F*(2,22) = 18.266, *p* < 0.001, η^2^ = 0.624) and instruction (*F*(1,11) = 26.311, *p* < 0.001, η^2^ = 0.705), and an interaction effect (*F*(2,22) = 18.266, *p* = 0.014, η^2^ = 0.322) on peak-CS (**Figures 5a, 6a**). As can be seen in **Figure 6a**, there were significant location effects during *fixation* (*F*(2,22) = 9.356, *p* = 0.003, η^2^ = 0.460), as well as *saccade preparation* (*F*(2,22) = 18.266, *p* < 0.001, η^2^ = 0.704). During *fixation*, peak-CS was significantly higher for the horizontal (*t*(11) = 6.137, *p* < 0.001, *d* = 0.903) and for the lower vertical meridian (*t*(11) = 2.655, *p* = 0.025, *d* = 0.678) than the upper vertical meridian. With *saccade* preparation, peak-CS was still higher for the horizontal (*t*(11) = 8.486, *p* < 0.001, *d* = 2.118) and the lower vertical (*t*(11) = 3.797, *p* = 0.005, *d* = 1.089) than the upper vertical meridian, similar to during *fixation*. In addition, peak-CS became higher for the horizontal than the lower vertical meridian (*t*(11) = 3.096, *p* = 0.013, *d* = 1.034), unlike during *fixation*.

**Figure 6.**
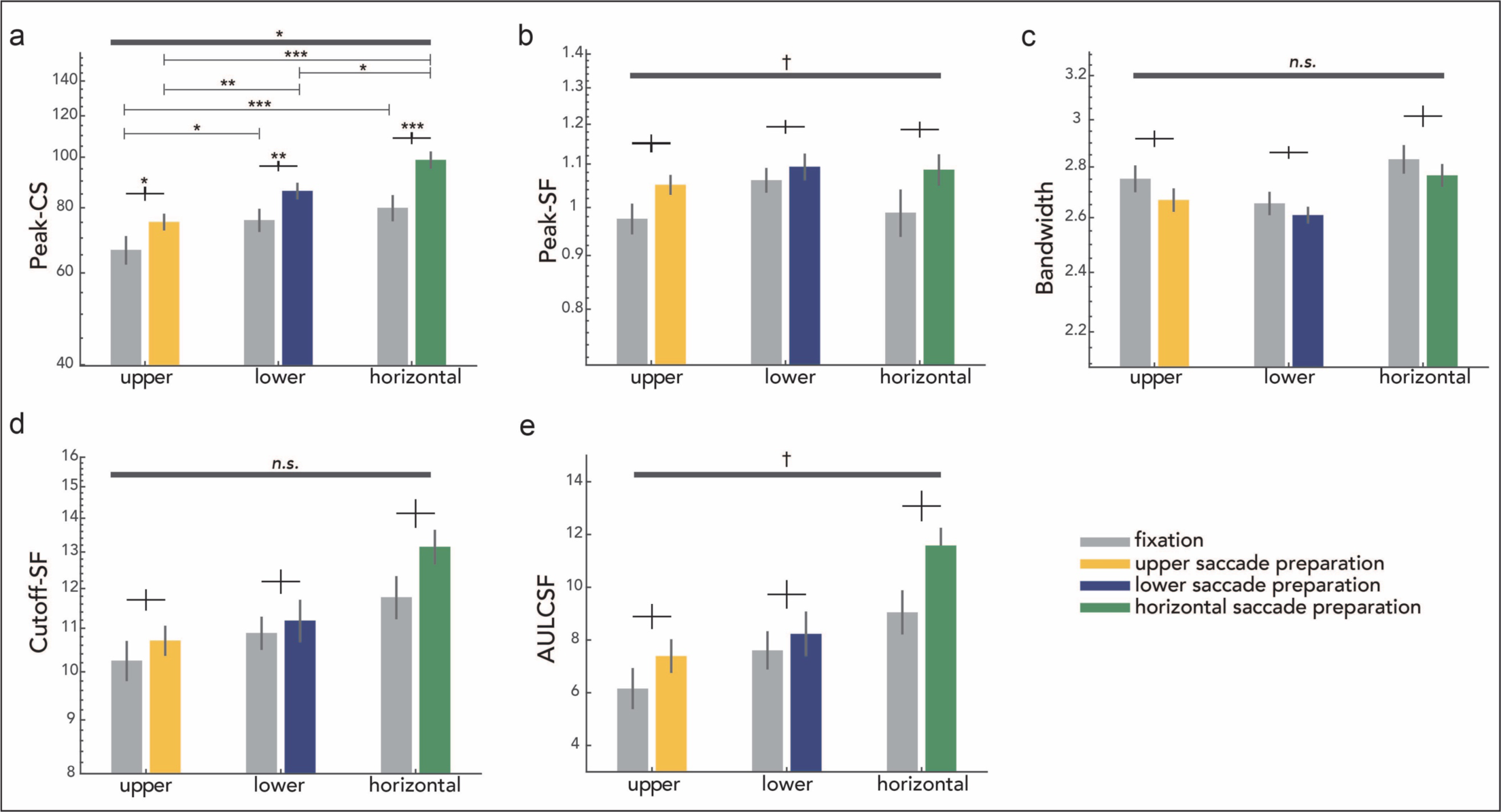
Group average key CSF attributes for each location and eye movement instruction. **(a)** peak-CS; **(b)** peak-SF; **(c)** bandwidth; **(d)** cutoff-SF; **(e)** AULCSF. Errorbars indicate ±_1_ standard error of the mean (SEM), and vertical bars depict ±_1_ standard error of the difference (SED). Location-instruction interaction effects are shown at the top of each figure with dark gray horizontal bars. Significant post-hoc pairwise comparisons of location and instruction effects are denoted by asterisks above black crosses and horizontal lines, respectively. \*\*\**p* < 0.001, \*\**p* < 0.01, \**p* < 0.05, †*p* < 0.1, *n.s*. not significant, FDR corrected.

*Saccade preparation* increased peak-CS compared to *fixation* for all locations (**Figure 6a**) J upper vertical (t(11) = 3.159, p = 0.013, d = 0.711), lower vertical (t(11) = 4.534, p = 0.002, d = 0.844), and horizontal (t(11) = 5.824, p < 0.001, d = 1.245) meridian, but the effect was larger for the horizontal meridian than the upper (t(11) = 2.833, p = 0.049, d = 0.158) and lower (t(11) = 2.370, p = 0.056, d = 0.151, BF_10_ = 2.083) vertical meridian. In summary, when compared to *fixation*, *saccade preparation* shifts the CSF upward and increases contrast sensitivity, for all locations, but more so for the horizontal meridian.

### 3.2. Peak-SF: spatial frequency leading to the peak-CS

There was a main effect of instruction (*F*(1,11) = 16.355, *p* = 0.001, η^2^ = 0.598), but not a main effect of location (*F*(2,22) = 1.756, *p* = 0.196, η^2^ = 0.138, BF_10_ = 1.454) on peak-SF (**Figure 5b**). The location-instruction interaction was marginally significant (*F*(2,22) = 3.303, *p* = 0.057, η^2^ = 0.231, BF_10_ = 0.335), resulting from presaccadic enhancement (increased peak-SF during *saccade preparation* relative to *fixation*) surviving FDR correction only for the horizontal (*t* = 3.082, *p* = 0.031, *d* = 0.728) and the upper vertical (*t* = 3.082, *p* = 0.031, *d* = 0.728) meridian (**Figure 6b**). Taken together, (1) peak-SF during *fixation* did not differ across locations; (2) *saccade preparation* increased the peak-SF compared to *fixation*, resulting in a rightward shift of the CSF; (3) the effect of *saccade preparation* on peak-SF was present at all locations. These results show that during *saccade preparation* contrast sensitivity is enhanced around the visual field particularly for spatial frequencies above the *fixation* peak-SF.

### 3.3 Bandwidth: full width at half maximum

There were main effects of location (*F*(2,22) = 7.137, *p* < 0.001, η^2^ = 0.396) and instruction (*F*(1,11) = 10.258, *p* = 0.005, η^2^ = 0.483, BF_10_ = 2.648), and no significant interaction (*F*(2,22) = 0.289, *p* = 0.752, η^2^ = 0.026, BF_10_ = 0.220) on bandwidth (**Figures 5c, 6c**). To further examine the main effect of location, bandwidth values were averaged across *fixation* and *saccade preparation* within each location for comparison across location pairs (**Figure 5c).** Only the difference between the lower vertical and the horizontal meridian survived FDR-correction: The bandwidth at the lower vertical meridian was narrower than at the horizontal meridian (*t*(11) = 4.142, *p* = 0.005, *d* = 1.134). The main effect of instruction reflects the bandwidth narrowing with *saccade preparation* (**Figure 5c**), and the lack of location-instruction interaction indicates that this effect is similar around the visual field (Figure 5c).

### 3.4. Cutoff-SF: highest perceivable spatial frequency at which contrast sensitivity is 1.0

We observed main effects of location (*F*(2,22) = 22.232, *p* < 0.001, η^2^ = 0.669) and instruction (*F*(1,11) = 18.250, *p* = 0.002, η^2^ = 0.624), and no significant location-instruction interaction (*F*(2,22) = 2.285, *p* = 0.125, η^2^ = 0.172, BF_10_ = 0.860) on cutoff-SF (**Figures 5d, 6d**). To follow up the location effect, we conducted post-hoc comparisons for location pairs on cutoff-SF values averaged across *fixation* and *saccade preparation* for each location (**Figure 5d**). The cutoff-SF at the horizontal meridian was significantly higher than that at the upper (*t*(11) = 7.787, *p* < 0.001, *d* = 1.324) and the lower vertical meridian (*t*(11) = 4.181, *p* = 0.002, *d* = 0.902). The main effect of instruction was caused by *saccade preparation* increasing the cutoff-SF (Figure 5d). This presaccadic enhancement on cutoff-SF was similar around the visual field (**Figure 6d**). These results, together with the increase in peak-SF, indicate that *saccade preparation* shifts the CSF rightward and enhances contrast sensitivity more for spatial frequencies higher than peak-SF during *fixation*.

### 3.5. AULCSF: area under the log CSF

We observed main effects of location (*F*(2,22) = 21.626, *p* < 0.001, η^2^ = 0.663) and instruction (*F*(1,11) = 26.642, *p* < 0.001, η^2^ = 0.708) on AULCSF (**Figure 5e**). The location-instruction interaction was marginally significant (*F*(2,22) = 3.109, *p* = 0.070, η^2^ = 0.220, BF_10_ = 0.207), from presaccadic enhancements in AULCSF surviving FDR correction only at the horizontal (*t* = 4.365, *p* = 0.003, *d* = 0.961) and the upper vertical (*t* = 2.602, *p* = 0.040, *d* = 0.500) meridian (**Figure 6e**). Post-hoc analyses of location effects were conducted on AULCSF values averaged across *fixation* and *saccade preparation* for each location (**Figure 5e**): AULCSF was significantly larger for the horizontal than the upper (*t*(11) = 8.725, *p* < 0.001, *d* = 1.486) and lower (*t*(11) = 3.900, *p* = 0.003, *d* = 0.957) vertical meridian. Thus, *saccade preparation* increases AULCSF at all locations, widening the window of visibility and rendering more of the world perceivable at the saccade target around the visual field.

### 3.6. Presaccadic benefit on contrast sensitivity across spatial frequencies

Saccade preparation improved performance across a wide range of spatial frequencies. There were main effects of location (*F*(2,22) = 7.162, *p* = 0.003 η^2^ = 0.394) and spatial frequency (*F*(50,550) = 9.957, *p* < 0.001, η^2^ = 0.143), and a significant interaction effect (*F*(100,1100) = 1.841, *p* < 0.001, η^2^ = 0.672) on the magnitude of the presaccadic benefit. To better understand the interaction effect, we conducted a one-way repeated measures ANOVA with location as a factor for each spatial frequency and observed a significant location effect in a wide spatial frequency range (**Figure 7**, between 0.616 and 11.314cpd, *p* < 0.05). The presaccadic benefit at the horizontal meridian was greater than at the upper vertical meridian (between 0.536 and 12.126cpd, *p* < 0.05) and the lower vertical meridian (between 0.758 and 11.314cpd, *p* < 0.05). The presaccadic benefit did not differ between the upper and lower vertical meridian for any of the spatial frequencies tested. Moreover, there was a maximum benefit of *saccade preparation* (relative to *fixation*) on contrast sensitivity for spatial frequencies higher than the *fixation* peak-SF (**Figure 7**), which is consistent with presaccadic attention shifting the peak-SF to higher spatial frequencies (**Figures 5b, 6b**).

**Figure 7.**
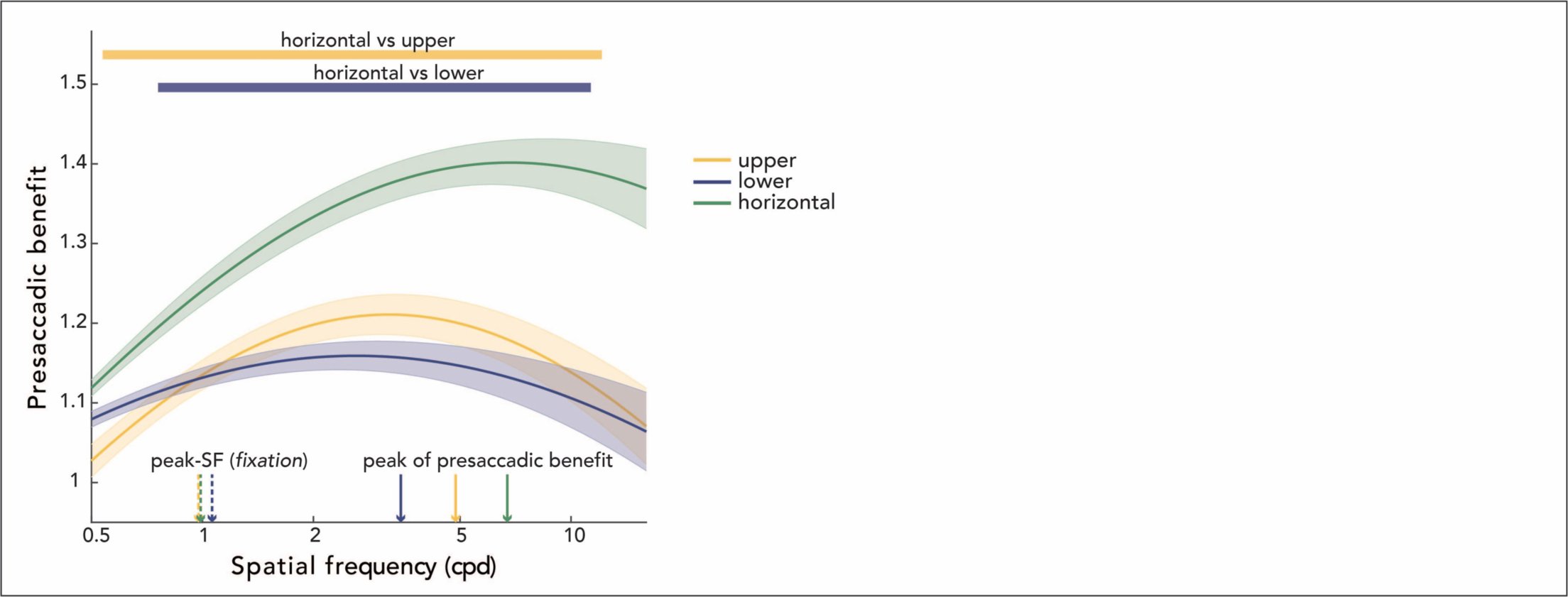
Presaccadic benefit, quantified as the ratio of contrast sensitivity during *saccade preparation* and *fixation*. Group average presaccadic benefit on the CSF, separately for each location. The yellow and blue horizontal bars at the top mark the spatial frequencies at which horizontal enhancement was larger than at the upper and lower vertical meridian, respectively. The vertical dashed arrows denote group peak-SF during *fixation* separately for each location. The vertical solid arrows indicate the spatial frequency resulting in the peak of the presaccadic benefit for each location. Note that the vertical solid lines do not necessarily correspond to the spatial frequency resulting in the maximum value of the group averaged curve shown in the figure. Shaded regions represent ±_1_ standard error of the mean (SEM).

### 3.7. Eye movement parameters

We evaluated eye movement parameters as a function of saccade direction, to determine whether these parameters influence presaccadic attention effects at different locations (**Figure 8**). A one-way repeated measures ANOVA yielded a main effect of saccade direction on latency, i.e., saccade onset relative to cue onset (*F*(1,11) = 22.459, *p* < 0.001, η^2^ = 0.672). Downward saccades were the slowest (mean = 228.015ms; **Figure 8a**), followed by horizontal (mean = 217.178ms) than upward (mean = 209.159ms) saccades (upward-downward: *t*(11) = 5.902, *p* < 0.001, *d* = 0.909; horizontal-downward: *t*(11) = 3.709, *p* = 0.002, *d* = 0.490; upward-horizontal *t*(11) = 3.524, *p* = 0.006, *d* = 0.419). There was neither a main effect of saccade direction on saccade amplitude (**Figure 8b**; BF_10_ = 1.126) nor on precision Dmean of the Euclidean distance between saccade endpoints and saccade target center (**Figure 8c**; BF_10_ = 0.458). Note that a minor difference in saccade latencies around the visual field has been reported previously (Honda & Findlay, 1992; Kwak et al., 2023; Schlykowa, Hoffmann, Bremmer, Thiele, & Ehrenstein, 1996; Tzelepi, Laskaris, Amditis, & Kapoula, 2010). The differences in saccade parameters do not parallel the perceptual differences and presaccadic attentional benefits around polar angle, and thus saccade parameters do not account for the more pronounced benefits in contrast sensitivity at the horizontal meridian.

**Figure 8.**
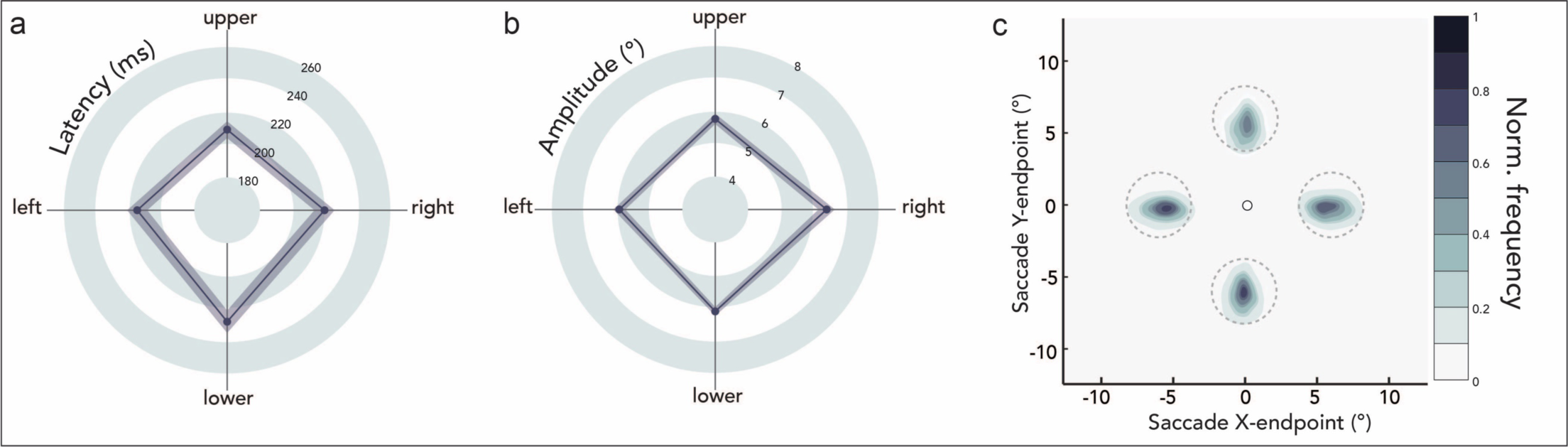
Group average saccade parameters as a function of saccade direction. **(a)** Saccade latencies (ms); **(b)** Saccade amplitudes (°). Shaded error areas indicate ±_1_ standard error of the mean (SEM). **(c)** Normalized saccade endpoint frequency maps averaged across participants as a function of saccade direction. Dashed gray circles indicate the 2.25° radius saccade target region. The central black circle indicates the fixation location (not on scale).

## 4. Discussion

This study reveals that presaccadic attention enhances and reshapes the contrast sensitivity function (CSF), and that the magnitude of these effects varies around the visual field. We compared key attributes of the CSF, estimated through Hierarchical Bayesian Modeling (HBM), during fixation and saccade preparation for different polar angle locations. Presaccadic attention increased the peak-CS, shifted the peak-SF and cutoff-SF to higher spatial frequencies, and sharpened the bandwidth (**Figure 9**), resulting in a larger area under the log CSF (AULCSF). Moreover, the increase in contrast sensitivity across a wide range of spatial frequencies, including the peak-CS, was more pronounced at the horizontal than the vertical meridian. Thus, during saccade preparation, the polar angle asymmetries in contrast sensitivity –specifically, the HVA (Horizontal-Vertical Anisotropy)– are exacerbated.

**Figure 9.**
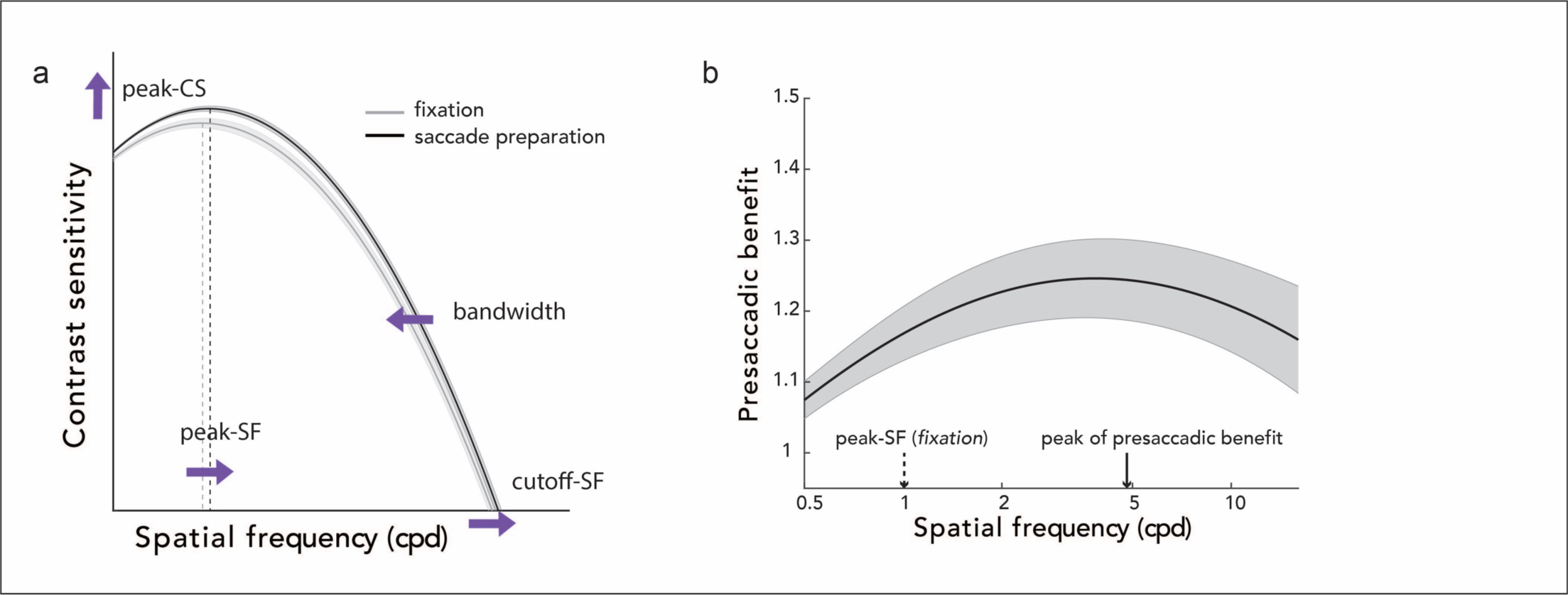
Summary schematic. (a) Key CSF attributes during fixation and saccade preparation. Presaccadic attention modulates peak contrast sensitivity (peak-CS), peak spatial frequency (peak-SF), cutoff spatial frequency (cutoff-SF), and bandwidth, which result in widening the window of visibility at the saccade target. Purple arrows demonstrate the direction of change. (b) Presaccadic benefit on contrast sensitivity across spatial frequencies. The benefit peaks at a spatial frequency (vertical solid arrow) higher than peak-SF during *fixation* (vertical dashed arrow). Shaded regions represent ±1 standard error of the mean (SEM).

These findings indicate that presaccadic attention renders the peripheral representation at the saccade target more fovea-like, in anticipation of the corresponding post-saccadic (foveal) input, which may facilitate a smooth transition in retinal images across saccadic eye movements (Herwig, 2015; Herwig & Schneider, 2014; Kroell & Rolfs, 2022; Li, Hanning, & Carrasco, 2021; Stewart, Valsecchi, & Schütz, 2020). Increases in peak-CS, peak-SF, and cutoff-SF, and sharpening of bandwidth during saccade preparation resemble the way the CSF changes from peripheral to foveal locations. Peak-CS and peak-SF gradually increase from the periphery to the fovea (Hilz & Cavonius, 1974; Jigo & Carrasco, 2020; Rovamo & Virsu, 1979; Rovamo et al., 1978). Moreover, cutoff-SF (i.e., acuity) is directly proportional to the cortical magnification factor and increases from peripheral to foveal locations (Cowey & Rolls, 1974; Rovamo & Virsu, 1979; Rovamo et al., 1978; Virsu & Rovamo, 1979). Further, the CSF bandwidth in visual cortex decreases from periphery to fovea (Goulet & Farivar, 2024).

Our findings are likely the perceptual consequences of neuronal modulations around the time of saccades. First, relevant to the role of presaccadic attention in retinal stability, neurons predictively shift, or ‘remap’, their receptive fields (RFs, to the location they will occupy post-saccade (Duhamel. Colby. & Goldberg, 1992: Neupane. Guitton, & Pack. 2016; Umeno & Goldberg, 1997). Accordingly, peripheral stimuli at the saccade target, such as the test stimulus in our study, would have been processed by neurons more selective for foveal locations -with smaller RFs. higher SF tuning. 3nd higher contrast sensitivity-during saccade preparation. Moreover. RFs shrink and shift toward the saccade target location, irrespective of the post-saccadic RF location (Neupane et al.. 2016; Tolias et al.. 2001). This leads to an increased proportion of neurons processing the saccade target. Both types of RF shifts predict the CSF parameter changes in the present study. Second, neuronal responses are enhanced, before saccade onset, when their RFs cover the saccade target location (Boch & Goldberg. 1989; Moore, Tolias. & Schiller, 1998). This mechanism can be particularly related to the observed presaccadic enhancement of peak-CS rather than of other CSF parameters in the SF space which likely rely on above-mentioned remapping mechanisms. Future studies should investigate whether distinct neuronal modulations underlie different perceptual changes during saccade preparation.

Our study reveals that the benefit of presaccadic attention on the CSF changes as a function of polar angle. During fixation, there were robust horizontal-vertical and vertical meridian asymmetries (HVA and VMA) effects in peak-CS. cutoff-SF. and AULCSF, but not in peak-SF. consistent with previous studies (Barbot, Xue, & Carrasco. 2021: Jigo et al.. 2023). During saccade preparation. the extent of the asymmetries for some of the CSF parameters was preserved; but for pe3k-CS as well as for a wide SF range, contrast sensitivity benefited more from saccade preparation at the horizontal than the vertical meridian, resulting in an even more pronounced HVA. Recent studies examining the interaction between presaccadic attention and polar angle asymmetries have yielded different results for contrast sensitivity (less benefit preceding upward saccades: Hanning et al.. 2022. 2023) and acuity (similar benefits around the visual field; Kwak et al.. 2023). The current study also uncovers a dissociation between location - instruction (fixation vs saccade preparation) interactions In peak-CS and cutoff-SF, which relate to contrast sensitivity and acuity measurements respectively, within the same experimental design and participants: Asymmetries in peak-CS as well as contrast sensitivity at a wide SF range were exacerbated with presaccadic attention, expanding contrast sensitivity more across the horizontal than the vertical meridian, whereas those in cutoff-SF remained. Hence, the interaction between presaccadic benefits and polar angle asymmetries may depend on the visual dimension. Importantly, polar angle asymmetries in both dimensions (i.e., contrast, SF) are not alleviated even by the deployment of presaccadic attention: They are either intensified (Hanning et al.. 2022. 2023) or preserved (Kwak et al.. 2023). Similarly, polar angle asymmetries are neither alleviated with covert spatial attention (Cameron et al.. 2002: Purokayastha. Roberts. Carrasco, 2021; Roberts, Cymerman, Smith, Kiorpes, & Carrasco, 2016) or temporal 478 attention (Fernandez, 479 Denison, & Carrasco, 2019).

What might cause the larger presaccadic benefit in the horizontal than the vertical meridian? Our results suggest that horizontal and vertical saccades might recruit different neuronal populations during saccade preparation. In fact, there is evidence that a neuron’s RF shifts toward the saccade target particularly for saccade directions aligned to the vector starting at fixation and ending at its RF center (Tolias et al., 2001). Therefore, it is possible that the presaccadic enhancement in the case of horizontal saccades arises from contribution of neurons with RFs along the horizontal, known to be more sensitive to various dimensions, than neurons along the vertical meridian, resulting in a larger benefit. Although this account does not explain the dissociation between presaccadic enhancement in contrast sensitivity and acuity, our findings call for investigation of the neuronal populations active for different saccade directions and visual dimensions.

As both the CSF and the benefit of presaccadic attention vary as a function of polar angle, our study alongside previous reports (Hanning et al., 2022, 2023; Jigo et al., 2023) call into question the generalizability of behavioral and neurophysiological measurements obtained along the horizontal meridian, the most frequently assessed to characterize the effects of target eccentricity as well as saccadic eye movements. Future studies should test whether oculomotor feedback signals (between eye movement and visual areas), which are thought to underlie presaccadic modulations of visual perception (Bisley & Mirpour, 2019; Ekstrom, Roelfsema, Arsenault, Bonmassar, & Vanduffel, 2008; Hanning, Fernández, et al., 2023; Moore & Armstrong, 2003), differ between horizontal and vertical saccades, and whether other perceptual correlates of presaccadic attention –such as the sensitivity shift to higher SFs (Kroell & Rolfs, 2021; Li et al., 2016, 2019) or orientation tuning (Li et al., 2016; Ohl et al., 2017)– are more pronounced for horizontal than vertical saccades.

In this study, we have *jointly* examined key attributes of the CSF in a systematic manner, across the whole contrast range, a wide range of SFs, and critically at different polar angle locations. Our findings conceptually correspond to presaccadic attention effects on *individual* attributes of the CSF tested *separately* in various visual tasks and dimensions. Presaccadic attention enhances contrast sensitivity at the saccade target when measured at a fixed SF (Hanning et al., 2022; Li, Pan, et al., 2021). It also enhances sensitivity to SFs higher than the stimulus SF (Li et al., 2016, 2019), but this was tested only for a narrow frequency range (1-2cpd) at 10° eccentricity along the horizontal meridian. These results are in line with the rightward shift of peak-SF and cutoff-SF reported in the present study. Moreover, the increased cutoff-SF is consistent with presaccadic attention enhancing visual acuity (Kwak et al., 2023). Lastly, for a constant contrast level, saccade preparation benefits orientation discrimination at horizontal saccade targets primarily at mid-SFs (Kroell & Rolfs, 2021). Here, measuring the CSF, we show that presaccadic benefit on contrast sensitivity peaks at SFs higher than the peak-SF during fixation around the visual field.

Deploying covert attention without concurrent eye movements 3lso modulates the CSF (Jigo & Carrasco. 2020). Endogenous attention, a voluntary and flexible mechanism, enhances contrast sensitivity across a broad range of SFs. Exogenous attention, an involuntary and inflexible mechanism, enhances contrast sensitivity primarily for SFs higher than the peak-SF at each eccentricity. The presaccadic attention benefits in the current study are reminiscent of both types of covert attention. As for endogenous attention, contrast sensitivity benefits are present across a wide range of spatial frequencies; at the peak SF during fixation, as well as below and above it As for exogenous attention, contrast sensitivity benefits are more pronounced for higher SFs ranges during saccade preparation, with peak-SF and cutoff-SF shifting to higher frequencies.

In conclusion, our comparison of the CSF at fixation and during saccade preparation has revealed that presaccadic attention not only enhances but also reshapes our contrast sensitivity across different SFs at the saccade target. Counterintuitively, presaccadic attention enhances our window of visibility more effectively at the horizontal meridian locations, where vision is typically stronger. This finding suggests that the integration of pre- and post-saccadic visual representations by presaccadic attention results in a more efficient smoothing of vision with horizontal than vertical eye movements.

## Acknowledgments

We thank members of the Carrasco Lab for helpful discussion and comments.

## Data availability statement

All materials will be available on the Open Science Framework database (https://osf.io/m8ysa) upon publication.

## Supporting Information

### Modeling prior distributions for HBM

For fitting the HBM, we start with the prior distributions of μ (population mean), Σ^,-1^ (inverse of population level covariance matrix), and φ^,!^ (inverse of individual level covariance matrix), which are *p*_1_(μ), *p*_0_(Σ^-1^), and *p*_1_(φ^,-1^) respectively.

For each of the CSF parameter, the prior distribution of μ is a truncated normal distribution: *N*(mean, standard deviation)T(lower bound, upper bound). The mean and standard deviation of the truncated normal distribution is set to the average of the mean (μ_*peakCS*,0_, μ_*peakSF*,0_, μ_*bandwidh*,0_) and standard deviation (σ_*peakCS*,0_, σ_*peakSF*,0_, σ_*bandwidth*,0_) of the corresponding parameters obtained from the BIP across individuals (*I*=72; combination of a participant, location, and instruction):

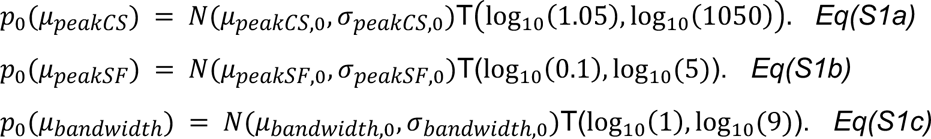

The prior distributions of the precision matrices (the inverse of covariance matrices Σ^−1^ and φ^−1^) are modeled as Wishart distributions. *W*(*Y, v*) denotes a Wishart distribution with expected precision matrix Y and degrees of freedom *v* (*v* = 4). Σ_*BIP*_ and φ_*BIP*_ are population-level and individual-level covariance matrices obtained from the BIP:

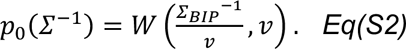

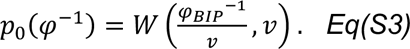

